# A statistical method for joint estimation of *cis*-eQTLs and parent-of-orign effects using an orthogonal framework with RNA-seq data

**DOI:** 10.1101/732792

**Authors:** Shirong Deng, Feifei Xiao

## Abstract

In the past few years extensive studies have been put on the analysis of genome function, especially on expression quantitative trait loci (eQTL) which offered promise for characterization of the functional sequencing variation and for the understanding of the basic processes of gene regulation. However, most studies of eQTL mapping have not implemented models that allow for the non-equivalence of parental alleles as so-called parent-of-origin effects (POEs); thus, the number and effects of imprinted genes remain important open questions. Imprinting is a type of POE that the expression of certain genes depends on their allelic parent-of-origin which are important contributors to phenotypic variations, such as diabetes and many cancer types. Besides, multi-collinearity is an important issue arising from modeling multiple genetic effects. To address these challenges, we proposed a statistical framework to test the main allelic effects of the candidate eQTLs along with the POE with an orthogonal model for RNA sequencing (RNA-seq) data. Using simulations, we demonstrated the desirable power and Type I error of the orthogonal model which also achieved accurate estimation of the genetic effects and over-dispersion of the RNA-seq data. These methods were applied to an existing HapMap project trio dataset to validate the reported imprinted genes and to discovery novel imprinted genes. Using the orthogonal method, we validated existing imprinting genes and discovered two novel imprinting genes with significant dominance effect.

**Author Summary:** In the past decades, an unprecedented wealth of knowledge has been accumulated for understanding variations in human DNA level. However, this DNA-level knowledge has not been sufficiently translated to understanding the mechanisms of human diseases. Gene expression quantitative trait locus (eQTL) mapping is one of the most promising approaches to fill this gap, which aims to explore the genetic basis of gene expression. Genomic imprinting is an important epigenetic phenomenon which is an important contributor to phenotypic variation in human complex diseases and may explain some of the “hidden” heritable variability. Many imprinting genes are known to play important roles in human complex diseases such as diabetes, breast cancer and obesity. However, traditional eQTL mapping approaches does not allow for the detection of imprinting which is usually involved in gene expression imbalance. In this study, we have for the first time demonstrated the orthogonal statistical model can be applied to eQTL mapping for RNA sequencing (RNA-seq) data. We showed by simulated and real data that the orthogonal model outperformed the usual functional model for detecting main effects in most cases, which addressed the issue of confounding between the dominance and additive effects. Application of the statistical model to the HapMap data resulted in discovery of some potential eQTLs with imprinting effects and dominance effects on expression of *RB1* and *IGF1R* genes.

In summary, we developed a comprehensive framework for modeling imprinting effect for eQTL mapping, by decomposing the effects to multiple genetic components. This study is providing new insights into statistical modeling of eQTL mapping with RNA-seq data which allows for uncorrelated parameter estimation of genetic effects, covariates and over-dispersion parameter.

## Introduction

With the completion the 1000 Genomes Project [1], an unprecedented wealth of knowledge has been accumulated for understanding variations in human DNA level. However, much less of this DNA-level knowledge has been translated to understanding the mechanisms of human diseases. Gene expression quantitative trait locus (eQTL) mapping is one of the most promising approaches to fill this gap, which aims to explore the genetic basis of gene expression [2]. Among the eQTL techniques, *cis*-eQTL mapping is the most commonly used technique to map local eQTLs on the same chromosome of the gene. To evaluate gene expression levels, RNA sequencing (RNA-seq) technology has recently become a widely used high-throughput tool to assess the gene expression abundance, which has many advantages over microarray, especially for discovery of novel eQTLs.

Imprinting is a type of parent-of-origin effect that the expression of certain genes depends on their allelic parent-of-origin. The same alleles transmitted from the mother may have different expression levels on transcripts with those transmitted from the father. The influence on the phenotype between the two types of heterozygotes are different, as so-called parent-of-origin effect (POE). It is now known that there are at least 80 imprinted genes in humans, many of which are involved in embryonic and placental growth and development [3]. Studies suggested that POE is an important contributor to phenotypic variation in human complex diseases and may explain some of the “hidden” heritability. An earlier study showed that for type II diabetes, a variant of SNP rs2334499 in chromosome region 11p15 was protective when maternally transmitted, whereas confers risk when paternally transmitted [4]. Evidence also demonstrated the important roles of POEs in type I diabetes, breast cancer and other carcinomas [4, 5]. However, most studies of eQTL maping have not implemented models that allow for the non-equivalence of parental alleles; hence the number and effects of imprinted genes remain important open questions. In the past few years, there were a few approaches that modeled POEs while searching for eQTLs with RNA-seq data. For example, Zhabotynsky et al. proposed to jointly model genetic effect and POE with a RNA-seq statistical method which focused on modeling the allele specific expression [6].

In quantitative genetics, the partition of the variance in statistical components due to additivity, dominance does not reflect the biological (or functional) effect of the genes but it is most useful for prediction, selection, and evolution [7]. In 2013, we proposed a unified orthogonal framework for modeling genetic variants displaying imprinting effects [8]. It allowed for imprinting effect detection whereas maintained the power to detect the main allelic effect including the additive and dominance effects. From theory of quantitative genetics, statistical additive genetic effects are obtained from average allele substitution effects, whereas dominance effects reflect the deviation of the genotypic values of the heterozygotes and the expected midpoint of the two homozygotes. In recent years, the inclusion of dominance in genomic models have been proposed by several researchers. However, multi-collinearity is an important issue arising from modeling multiple genetic effects. To achieve straightforward model selection and variance component analysis, uncorrelated estimation of the additive and dominance effect was necessary. Therefore, in current study, we propose to implement an orthogonal model to jointly evaluate the effect from both additive and dominance effect along with POE in eQTL mapping.

Genetic imprinting affects complex diseases through regulating the gene expression and can reveal an important component of heritable variation that remains “hidden” in traditional complex trait studies. In this study, we hypothesized that POEs contribute to regulate gene expression along with the main allelic effect from the gene. We proposed a statistical framework to test the main allelic effects (i.e., additive and dominance effects) of the candidate eQTLs along with the POE with an orthogonal model. Intensive simulations were conducted to evaluate the methods. We also applied the methods to an existing HapMap project trio dataset to validate the reported imprinting genes and identifying eQTLs for these genes.

## Results

### Simulations

The statistical power of the Stat-POE and Func-POE methods was illustrated when the POE was (a) τ = log(1.1) and (b) τ = log(1.2). The results are shown in Figures 1–2, respectively. In both scenarios, the additive and dominance effects were fixed at log (1.2). We simulated a small fold change in both allelic effect terms to demonstrate the desirable performance of the Stat-POE method when the effect size were relatively small to detect. Consequently, even at a sample size of 50 with moderate over-dispersion (φ= 0.2), the Stat-POE method presented around 70.8% power to detect a genetic effect at 1.2-fold change in additive effect, corresponding to an effect size of log(1.2) = 0.18 (Figure 1). To detect POE at the fold change of 1.2, the Stat-POE and Func-POE methods both reached a statistical power of 83% with a reasonable sample size of 100 (Figure 2). Even with a very small effect size from POE at fold change of 1.1, corresponding to an effect size of log (1.1) = 0.10, the methods yielded 61% power when the sample size was 200, and 91% when the sample size was 500 (Figure 1). As expected, the Stat-POE method yielded same power in detecting POE but more desirable power in detecting main genetic effects compared to the Func-POE model (Figures 1–2). In conclusion, the Stat-POE method outperformed the Func-POE method in most simulation scenarios and our methods all achieved sufficient power for detection of POEs with a relatively sample size (N=100).

**Figure 1.**
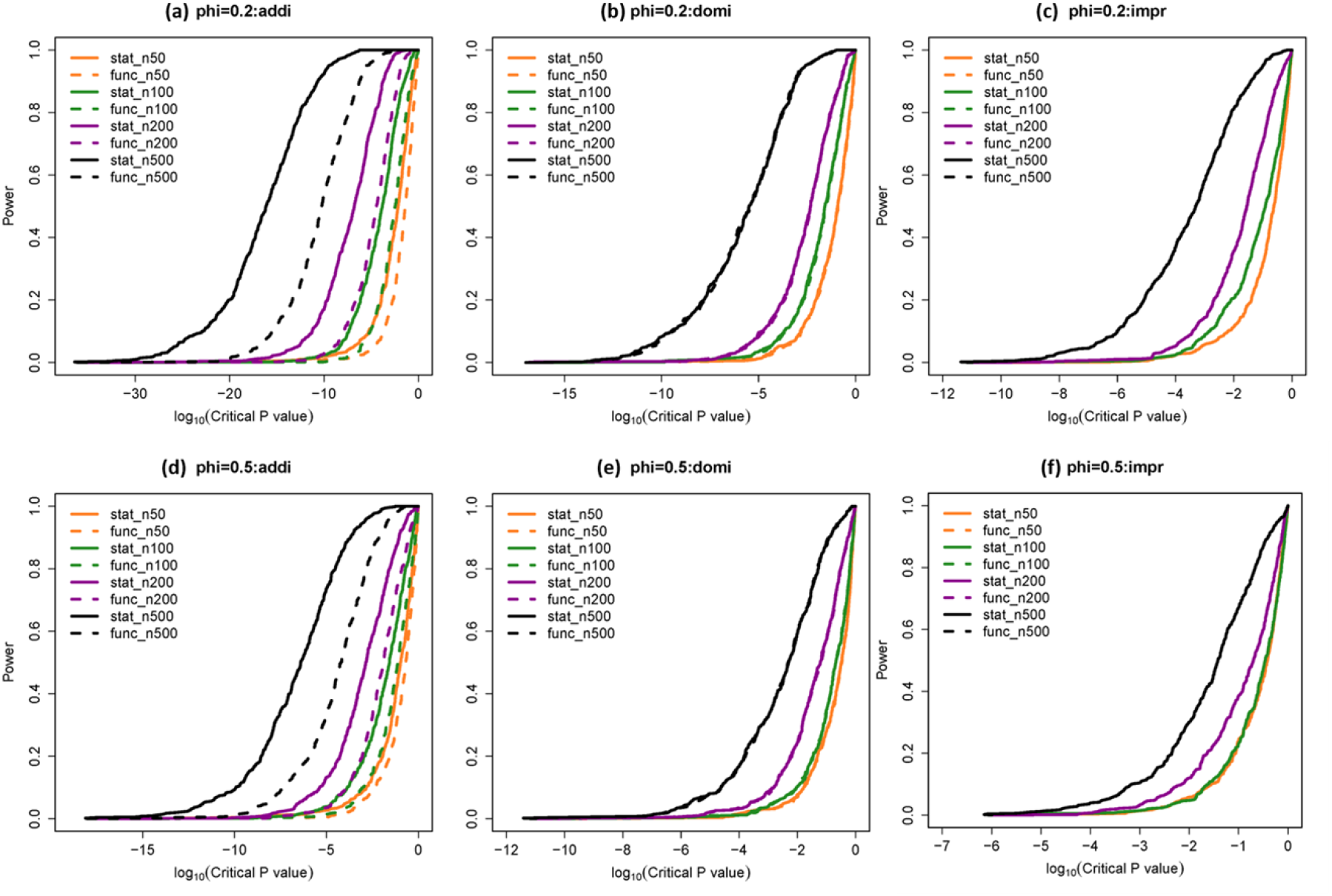
**Statistical power to detect additive, dominance and POE effect** when (a)-(c) overdispersion φ = 0.2 and (d)-(e) φ = 0.2 for various samples sizes, using Stat-POE model (stat) or Func-POE model (func). The covariate coefficient *β* = 0.1, the sample size (n) was set at 50, 100, 200 and 500. Addi: additive effect; domi: dominant effect; impr: imprinting effect; stat: statistical model; func: functional model. The additive effect α = log(1.2), dominant effect δ = log(1.2), imprinting effect ι = log(1.1).

**Figure 2.**
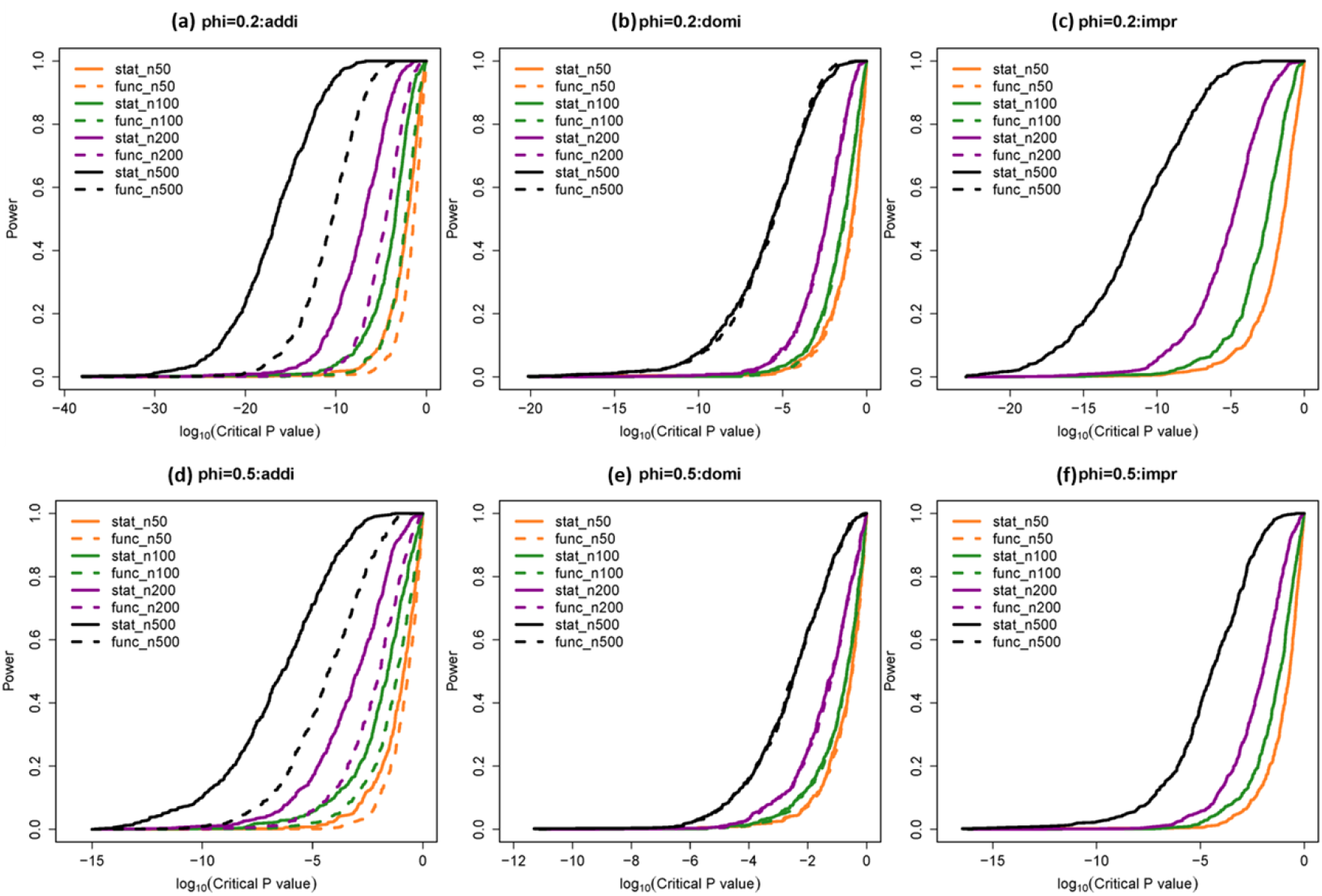
**Statistical power to detect additive, dominance and POE effect** when (a)-(c) overdispersion φ = 0.2 and (d)-(e) φ = 0.2 for various samples sizes, using Stat-POE model (stat) or Func-POE model (func). The covariate coefficient *β* = 0.1, the sample size (n) was set at 50, 100, 200 and 500. Addi: additive effect; domi: dominant effect; impr: imprinting effect; stat: statistical model; func: functional model. The additive effect α = log(1.2), dominant effect δ = log(1.2), imprinting effect ι = log(1.2).

With the simulated data, we also evaluated the estimation bias for all the parameters (*β,β_G,φ_*) from the Stat-POE model. Table 1 shows that the estimation of all genetic effects achieved higher accuracy when sample size increased. Interestingly, the estimation of the covariate *β* and over-dispersion parameter *φ* was not notably affected by the sample sizes. Also, the estimation of genetic effects was not obviously affected by the value of the over-dispersion parameter. These results revealed the accurate and stable estimation of the covariates and over-dispersion parameters by using the Stat-POE model. Moreover, large sample sizes and small overdispersion ensured better overall performance of the proposed methods.

**Table 1.** Simulation results with different sample sizes. The estimation relative bias was defined as the difference between estimated value and true parameter value divided by the true parameter value. Relative bias and mean square of errors (MSE) have been multiplied by 100. for model parameters including additive effect (α), imprinting effect (ι), covariate (β), and dispersion parameter (φ).

|  |  | N=200 |  |  |  |  |  | N=500 |  |  |  |  |  |
| --- | --- | --- | --- | --- | --- | --- | --- | --- | --- | --- | --- | --- | --- |
|  |  | Bias |  |  | MSE |  |  | Bias |  |  | MSE |  |  |
| Over-dispersion | True POE effect size | $\alpha$ | $\delta$ | $\iota$ | $\alpha$ | $\delta$ | $\iota$ | $\alpha$ | $\delta$ | $\iota$ | $\alpha$ | $\delta$ | $\iota$ |
| 0.2 | log(1.1) | -0.98 | 1.96 | -0.47 | 0.21 | 0.44 | 0.20 | 1.15 | -0.83 | 2.35 | 0.09 | 0.18 | 0.07 |
| 0.2 | log(1.2) | -1.80 | 1.67 | -1.12 | 0.21 | 0.39 | 0.22 | -0.18 | -0.03 | -0.41 | 0.09 | 0.18 | 0.08 |
| 0.5 | log(1.1) | -3.48 | -1.57 | -6.65 | 0.49 | 1.08 | 0.58 | -2.47 | 0.46 | -1.84 | 0.20 | 0.40 | 0.19 |
| 0.5 | log(1.2) | -0.71 | 2.13 | 0.66 | 0.52 | 1.04 | 0.52 | -0.34 | 0.85 | 0.07 | 0.22 | 0.42 | 0.17 |

To evaluate the Type I error, we also simulated scenarios where there no genetic effect or POE. Although there were slightly inflated false positives in detecting genetic effect and POEs when over-dispersion was large, the type I error rates were around the nominal level 0.05 for both methods in most scenarios especially when moderate over-dispersion existed (*φ* = 0.2 (Table 2). The false positive rate for detecting the genetic effects was comparable between the Stat-POE and Func-POE models in most scenarios.

**Table 2.**
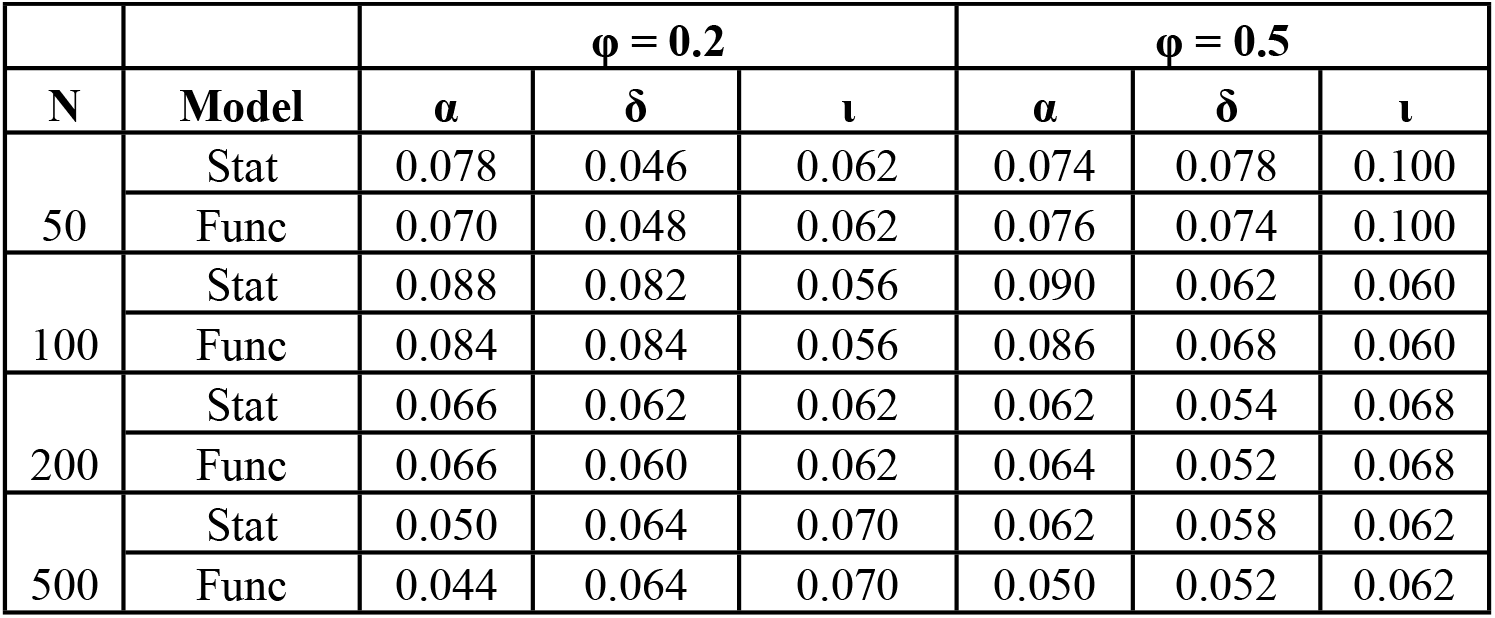
Type I error of the methods in detecting genetic effects. Stat: Stat-POE model; Func: Func-POE model. α: additive effect; δ: dominant effect; ι: imprinting effect, and φ: dispersion parameter

### Real Data Application to HapMap Parent-Child Trio Data

Using 30 children of the family trios from the HapMap project, we applied the proposed methods to estimate additive and imprinting effects for 22 genes with previous evidence of imprinting. These selected genes were identified to be imprinted genes using 296 phased trios from the 1000 Genomes Project and the Genome of the Netherlands participants [9].

With the proposed Stat-POE method, we identified 33 significant *cis*-eQTLs (adjusted p-values in additive effects < 0.05) for seven genes including *LPAR6, RB1, PXDC1, IGF1R, AC069277.2, IGF2BP3, SNRPN* genes (S2 Table). Among them, most candidate *cis*-eQTLs presented maternal expression pattern in regulating the gene expression. Besides, we identified six genes with imprinting effects, including *LPAR6, PER3, RB1, PXDC, IGF1R*, and *IGF2BP3* with adjusted p-values < 0.05 (Table 3). Among the significantly imprinted genes, the gene expression of *LPAR6* and *IGF1R* had significant regulation from the candidate *cis*-eQTLs rs11633209, rs728075 and rs7329291 in additive effect (adjusted p-values = 1.96×10^−66^, 3.57×10^−64^ and 3.57×10^−64^). Interestingly, we also discovered two novel genes which presented significant dominance effect in gene expression (Table 4), including *RB1* gene from multiple candidate *cis*-eQTLs (adjusted p-value = 3.02×10^−80^) and *IGF1R* gene from rs4965238 (adjusted p-value = 1.37×10^−67^). Among the identified genes presenting dominance effect in eQTL mapping, the *RB1* gene locating on chromosome 13 is a tumor suppressor gene, the mutation inactivation of which has been found to be the cause of human cancer [10]. It was also found to be an imprinted gene earlier in 2009 [11]. Interesting, *IGF1R* is the only gene which presented both dominance effect and imprinting effect from the candidate *cis*-eQTL rs4965238 (Table 4).

**Table 3.** A list of genes with potential imprinting effects (p-values < 0.05). ATL: alternative allele; a: additive effect; d: dominance effect; l: imprinting effect; expression: imprinting status. The p-values were adjusted by the BH multiple comparison method.

| ID | Gene | Chr | SNP | Position | Allele | ATL | P value.a | P value.d | P value.l | Expression |
| --- | --- | --- | --- | --- | --- | --- | --- | --- | --- | --- |
| ENSG00000049246 | *PER3* | chr1 | rs67537516 | 7864693 | G | A | 0.96 | 1 | **0** | Maternal |
| ENSG00000139687 | *RB1* | chr13 | rs61949059 | 48923973 | T | G | 0.18 | 1 | **6.26E-82** | Maternal |
| ENSG00000139687 | *RB1* | chr13 | rs150098624 | 49001371 | T | C | 0.9 | 1 | **3.06E-80** | Paternal |
| ENSG00000139687 | *RB1* | chr13 | rs4258502 | 48989564 | G | A | 0.9 | 1 | **3.06E-80** | Paternal |
| ENSG00000139687 | *RB1* | chr13 | rs9568143 | 48996494 | T | A | 0.9 | 1 | **3.06E-80** | Paternal |
| ENSG00000140443 | *IGF1R* | chr15 | rs4965238 | 99456357 | T | C | 0.78 | 1.37E-67 | **1.12E-71** | Paternal |
| ENSG00000140443 | *IGF1R* | chr15 | rs11633209 | 99221205 | A | T | **1.96E-66** | 1 | **5.01E-71** | Maternal |
| ENSG00000168994 | *PXDC1* | chr6 | rs113644426 | 3738592 | C | T | 0.98 | 1 | **4.14E-69** | Paternal |
| ENSG00000139679 | *LPAR6* | chr13 | rs4942796 | 49011882 | T | C | **0** | 1 | **5.81E-64** | Maternal |
| ENSG00000139679 | *LPAR6* | chr13 | rs728075 | 48999087 | G | A | **3.57E-64** | 1 | **5.81E-64** | Maternal |
| ENSG00000139679 | *LPAR6* | chr13 | rs7329291 | 48999148 | G | A | **3.57E-64** | 1 | **5.81E-64** | Maternal |
| ENSG00000139679 | *LPAR6* | chr13 | rs4942795 | 49003191 | A | G | **0** | 1 | **7.86E-64** | Maternal |
| ENSG00000139679 | *LPAR6* | chr13 | rs7998359 | 48983977 | G | A | **0** | 1 | **7.86E-64** | Maternal |
| ENSG00000140443 | *IGF1R* | chr15 | rs62023801 | 99295429 | T | C | 0.99 | 1 | **0.001** | Maternal |
| ENSG00000140443 | *IGF1R* | chr15 | rs6598236 | 99299001 | G | A | 1 | 1 | **0.001** | Maternal |
| ENSG00000136231 | *IGF2BP3* | chr7 | rs56259943 | 23400797 | C | T | 0.15 | 1 | **0.0137** | Maternal |
| ENSG00000136231 | *IGF2BP3* | chr7 | rs73073066 | 23447351 | C | T | 0.22 | 1 | **0.0207** | Maternal |

**Table 4.**
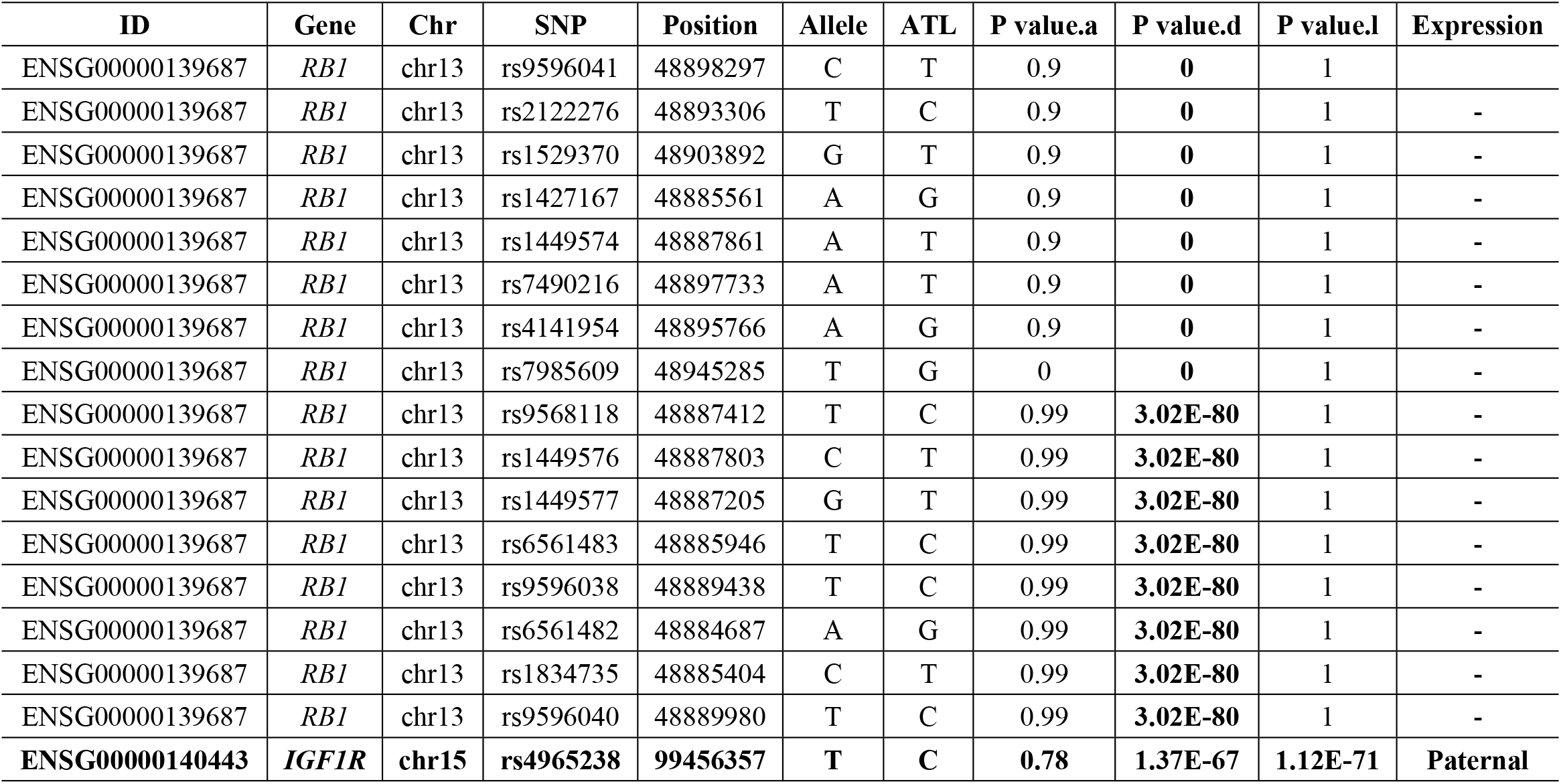
A list of genes with potential dominance effects (p-values < 0.05). ATL: alternative allele; a: additive effect; d: dominance effect; l: imprinting effect; expression: imprinting status. The p-values were adjusted by BH multiple comparison.

In conclusion, our real data application validated several existing imprinted genes, mapped candidate eQTLs for these imprinted genes. More interestingly, we discovered a few genes presenting significant dominance effect which might be involved in tumorigenesis.

## Discussion

This article stands on recent advances in genetic modeling for carrying out new methodological developments to the aid of the analysis of genetic imprinting. The orthogonal statistical model here developed improves previous methods by providing a solution to allowing for additive-by-dominance genetic effects for *cis*-eQTL mapping with RNA-seq data. We demonstrated the desirable power and preserved Type I error of the statistical POE model with un-biased estimation of the genetic effects and over-dispersion of the RNA-seq data. The application to the HapMap project validated previously reported imprinting genes and discovered significant *cis*-eQTLs for these imprinted genes. More interestingly, we also identified two novel imprinting genes with significant dominance effect, although the interpretation of which is still unclear.

This is the first time the performance of the natural and orthogonal models in detecting POE with RNA-seq data are evaluated. This project is an extension of the natural and orthogonal interaction model (NOIA) proposed by Albarez-Castro et al., in 2007 [12], based on which we have extensively worked on the framework to the estimation of statistical epistasis, gene-environmental interactions and imprinting effect in genotype-phenotype mapping for quantitative traits and qualitative traits [8, 13, 14]. In previous findings, we proved that the one-locus Stat-POE model for quantitative trait was orthogonal in certain conditions including when HWE was satisfied [8]. The conclusion should be straightforwardly extended to negative binomial or Poisson regressions. Although it still needs more effort to evaluate the orthogonality of the Stat-POE model when HWE is violated for the current framework of eQTL mapping in RNA-seq, the proposed work is providing new insights into eQTL mapping which allows uncorrelated parameter estimation of genetic effects (i.e., additive effect, dominance effect and imprinting effect) and accurate estimation of covariates and over-dispersion parameters.

In this study, we compared the Stat-POE model and Func-POE model for analyzing POE effects for RNA-seq data. These two models can be transformed to each other, but they had different meanings for their parameters and their test statistics varied in formulas. The Func-POE model focus on the biological properties and describes allele substitution effects thus presented a natural and interpretable results. The Stat-POE model utilizes the population properties estimated from the sample therefore presented greater power in detecting additive effect compared to the Func-POE model, which was consistent with our previous methodology studies in genotype-phenotype mapping for quantitative traits [8]. However, using the estimates of genotype frequencies, parameters from application of the Stat-POE model describes the variance components, rather than allele substitutions effects as in Func-POE model, so may be seen as having a less clear interpretation. The results generated from the functional model is not tied to population rendering easier interpretation however the model selection is not straightforward. In applications, we will suggest utilizing both Stat-POE and Func-POE models to further characterize the underlying genetic architecture of the eQTLs.

We also investigated the parameter estimation and hypothesis testing of the two models with NB regression and Poisson regression assumption of the read counts for different application scope (results not shown). NB regression will be suggested when the over-dispersion is relatively large and Poisson regression is suggested to be used for small over-dispersion. In conclusion, we provided a comprehensive and robust framework for joint estimation of *cis*-eQTLs and parent-of-origin effects.

The current study has several limitations. First, family data such as trios are needed to obtain the genotypes of heterozygotes in the offspring. Second, borrowing information from the whole samples will allow for better power for modeling the RNA-seq data. A third direction is to incorporate the allele specific gene expression(ASE) as conducted in Zhabotynsky’s work so we can extend our study to model both ASE and POE for candidate *cis*-eQTLs [6]. The gradually decreased cost in RNA-seq technology and future studies in methodology are warranted to achieve more powerful estimation of decomposed variance from different genetic components.

## Materials and Methods

### Methods: the Stat-POE and Func-POE methods

Our methods are proposed upon a basic model of eQTL mapping of a single gene with RNA-seq data which are read counts. Therefore, we consider a single gene and study the association of its expression with the *j*-th candidate expression quantitative trait loci (eQTL). Let *y_i_* be the total read counts mapped to this gene in the *i*-th sample, where *i* = 1,…,*n* and *n* is the sample size. We model *y_i_* by a negative binomial distribution as they are sparse counts data, which is a generalization of Poisson distribution to allow for over-dispersion. Let *f_NB_*(*y_i_*;*μ_i_*,*ϕ*) be the probability mass function for a negative binomial distribution with mean *μ_i_* and dispersion parameter *ϕ*:

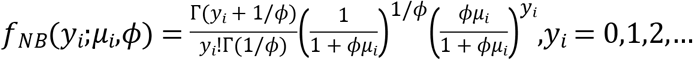

where Γ(·) is the gamma function. It’s easy to find that the variance 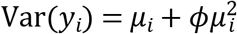, in which 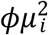 is the over-dispersion part. When the over-dispersion parameter *ϕ* goes to 0, *f_NB_*(*y_i_*;*μ_i_*,*ϕ*) reduces to 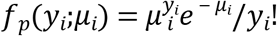, which is the probability mass function for Poisson distribution with parameter *μ_i_*. Let **X**_*i*_ be a set of p covariates and ***β*** = (*β*_1_,…,*β_p_*)′ be the regression coefficients, and ***β***_*G*_ = (*R,a,d,l*)′ be the genetic effects from genotypes (*G*) of the eQTL on *Y*, where *R* is the baseline, *a, d* and *l* are the additive, dominance and imprinting effects from *G*, respectively. The covariate effect of *G* = *G_i_* and covariates **X** = **x**_*i*_ on the gene expression, can be formulated through the following log-linear regression model

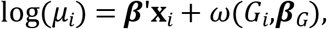

where *ω*(*G_i_*,***β***_*G*_) is the function which reflects the genetic effects.

For a bi-allelic locus, let the major and minor alleles of the *j*-th candidate eQTL as *A*_1_ and *A*_2_, respectively. The genotype *G* takes four possible values 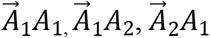 and 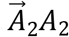, the first allele of which with arrow denotes the paternal allele and the second denotes the one originated from maternal side. We use *p*_11_,*p*_12_,*p*_21_ and *p*_22_ to denote genotype frequencies in the population, and use *M* to denote the number of variant allele *A*_2_, which takes values of 0, 1, 1 and 2 for the four genotypes separately. 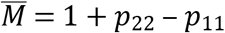 and *V* = (*p*_11_ + *p*_22_) – (*p*_11_ – *p*_22_)^2^ are the mean and variance of *M*. For estimation of the genetic effects, there are different methods we can epress the genetic effect function *ω*(*G_i_*,***β***_*G*_). Xiao (2013) proposed two related methods for identifying genetic variants influences quantitative traits with different characteristics, the statistical and functional POE estimation methods [8]. Accordingly, we can express *ω*(*G_i_*,***β***_*G*_) as a population-referenced formulation using an orthogonal model, such expression generated from orthogonality reparameterization:

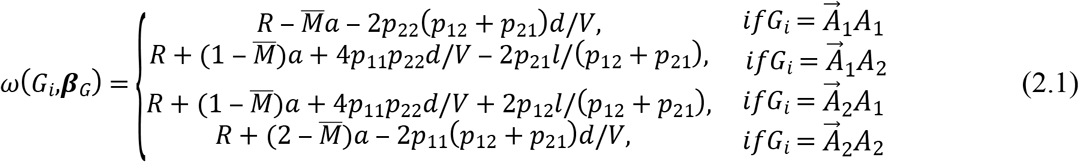

With the orthognality property, this model allows for un-correlated estimation of the genetic effects including additive, dominance and imprinting effect. Therefore it is referred to the statistical POE (Stat-POE) model. For a functional model with no orthogonalization property with a POE component,, the genetic effect function *ω*(*G_i_*,***β***_*G*_) can be expressed as

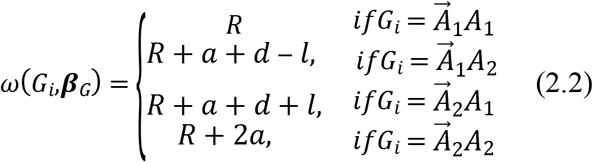

from which we obtain

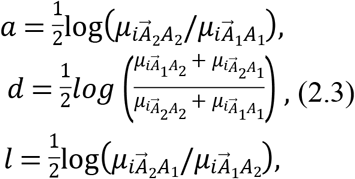

where 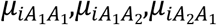 and 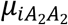 are the underlying means of the read counts for subjects with the four genotypes, respectively. Indeed, the additive effect *a* measures the fold change of gene expression between the two homozygotes; the dominance effect *d* measures the deviation of the heterozygotes from its additive expectation; and the imprinting effect *l* reflects the different effect from the two types of heterozygotes. The model in Equations (2.3) is therefore defined as a functional POE (Func-POE) model. This genetic effect model can also be called as natural model since it uses natural effects of allele substitutions as parameters, mainly focusing on the biological properties [12].

Theoretically, the parameters from the Stat-POE and Func-POE models can be transformed to each other in linear regression setting, yet which still present different properties for various application scopes. As shown in the theoretical derivation from Xiao (2013) [8], the statitical model will present better power in detecting additive effect given the existence of this effect whereas the functional model generates more interpretable results.

### Parameter Estimation and Hypothesis testing

To estimate the genetic effects and POE, we can write the likelihood based on the data (*y_i_,X_i_,G_i_*) (*i* = 1,2,…*N*) as follows,

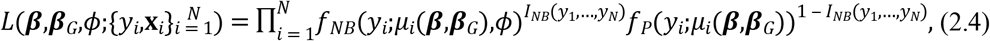

where *I_NB_*(*y*_1_,…,*y_N_*) is an indicator function which equals to 1 if a negative binomial distribution is used and 0 if a Poisson distribution is used.

With equation 2.4, the maximum likelihood estimator (MLE) of the model parameters (***β***_*ϕ*_,***β***_*G*_) with ***β***_*ϕ*_ ≜ (***β***′,*ϕ*)′ can be estimated by the following iterative procedure.

1. **Initialization:** We first fit a null model by a Poisson regression using the covariate **X**_*i*_, and then estimate ***β***, which is obtained by the Newton Raptson iteration method based on the first and second derivatives for the log-likelihood function of the null model, whose formulas are given in Appendix A.1. Then a score test is conducted for the over-dispersion parameter, the whole hypothesis testing process of which is shown in Appendix A.2. If the p-value of the score test is smaller than a cutoff value, e.g., *α* = 0.05, we will fit a negative binomial regression model and estimate ***β***_*ϕ*_. Under the negative binomial model, the estimate of ***β***_*ϕ*_ is obtained by an iterative procedure. The details of the iterative formulas for ***β*** and *ϕ* are given in (A.10) and (A.11) in Appendix A.3, which is based on the iteratively re-weighted least squares method [15] and the Newton-Raphson iterate method, respectively.
2. **Iteration**: (a). Given ***β*** or ***β***_*ϕ*_, estimate ***β***_*G*_ by the Newton-Raphson method, the details of which are shown in **Appendix B**; (b). Given ***β***_*G*_, estimate ***β*** by a Poisson regression with offsets *ω*(*G_i_*,***β***_*G*_), or estimate ***β***_*ϕ*_ by a negative binomial regression with offsets *ω*(*G_i_*,***β***_*G*_). The estimation for ***β*** under the Poisson regresion is the same as that in the initialization step with the first and second derivatives given in Appendix B.6 for the statistical model and Appendix B.8 for the functional model, respectively. Under the negative binomial regression, the estimation method for ***β***_*ϕ*_ described in the the initialization step is also used here with the detailed forlumas given in Appendix B.5 for the statistical model and in Appendix B.7 for the functional model respectively.
3. **Termination:** Iterate steps (1) and (2) until estimates of all the parameters converge.

In order to assess whether each regression covariate in the model is significant on the read counts of the gene or not, statistical hypothesis testing will be performed. We constructed three testing methods including the likelihood ratio test (LRT), score test and wald test as follows. For example, for the test of additive effect with hypotheses

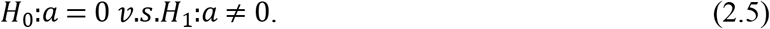

Denote 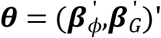 for the Negative Binomial (NB) regression, or 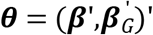 for the Poisson regression, the unrestricted MLE and restricted MLE under (Appendix D.1) obtained by the algorithm given in above section are denoted as 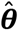 and 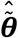 respectively. Without loss of generality, we put the parameter *a* in the first position of ***θ***, denote the other parameters as *ξ*, i.e. ***θ*** = (*a,ξ*′)′.

Then the score function for ***θ*** is 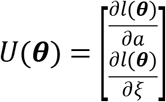, and the observed fisher information matrix is 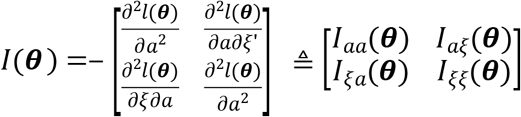, where *l*(***θ***) is the log-likelihood function given as in Appendix B.1 for NB regression and statistical model, in Appendix B.2 for Poisson regression and statistical model, in Appendix B.3 for NB regression and functional model, in Appendix B.4 for Poisson regression and functional model. The formulas of *U*(***θ***) and *I*(***θ***) are given in **Appendix C**. The LRT statistic is

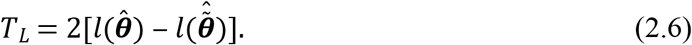

According to the theory from Rao 2005 [16], in our statistical setting, the score test statistic is therefore defined as

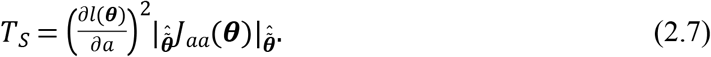

where 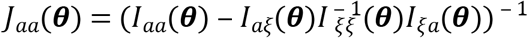.

Moreover, the Wald test statistic is defined by:

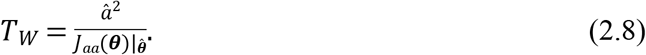

Under *H*_0_, the statistics *T_L_, T_S_*, and *T_W_* all converge in distribution to 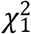. For a given significance level *α*, reject *H*_0_ when the observed value of the statistics are greater than 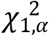. The process of the hypothesis testing for the other parameters can be implemented similarly.

### Simulations

To evaluate the performance of the proposed statistical methods in eQTL mapping with RNA-seq data, we carried out extensive simulation studies in realistic settings. First, we compared power of the Stat-POE and Func-POE methods in detecting the main allelic effects (including the additive and dominance effects) and POE. We simulated *y_i_*, the total number of read counts of a gene in the *i*-th sample by a negative binomial distribution with *μ_iG_* = exp(0.1*x_i_* + *ω*(*G_i_,β_G_*)) and over-dispersion parameter *φ* = 0.2, where **X** was a continuous covariate **X**~N(0,1). To evaluate the performance of the methods in estimating both genetic effects and over-dispersion, we generated data with different sample sizes N = 50, 100, 200 and 500, respectively. In all simulation settings, Hardy-Weinberg Equilibrium (HWE) was assumed that the genotype frequencies in the sample were set at [*p*_11_, *p*_12_, *p*_21_, *p*_22_] = [0.36, 0.24, 0.24, 0.16]. The overdispersion parameter was set at empirical values which was 0.2 or 0.5. The additive effect α and dominance effect δ were both fixed at log (1.2) where the values of 1.2 reflected the fold change of the logarithm mean shift of the genotypic values, referring to Equation (2.3). The POE parameter τ was set at log (1.1) or log (1.2) respectively. Each simulation was replicated for 500 times to evaluate the performance of the Stat-POE and Func-POE methods. Relative bias and mean square of errors (MSE) were calculated for each parameter in the different scenarios to evaluate the estimation accuracy. The estimation relative bias was defined as the difference between estimated value and the true parameter value and then divided by the true parameter value. We also used simulated data to quantify the power and Type I error rates of the methods. To illustrate the performance of the proposed methods in detecting different genetic effect terms and POE, the statistical power for the hypothesis testing was calculated using a range of different critical values under different scenarios. Type I error was calculated under the null model where there was no genetic effect or POE.

### Application to a HapMap RNA-seq dataset

#### Datasets

We used an RNA-seq dataset from 30 HapMap Caucasian samples obtained from the NCBI Bioproject (PRJNA385599). The samples were collected from lymphoblastoid cell lines from 15 males and 15 females. For most of these samples, the RNA reads were 150 bp paired-end reads, with an additional run with 75 bp paired-end reads. The median of the total number of reads for these 30 samples was approximately 20 million. All of these reads were mapped to hg38 human reference genome using Tophat2 [6].

Since all of these samples were children of family trios, the parents of which were also part of the samples included in the 1000 Genomes Project[17]. Genotype data of the 30 trios were used to obtain the phased genotype of the children. For these 30 trios, the HapMap project genotyped about 3.9 million SNPs. The phasing and imputation of these 30 trios were conducted by Zhabotynsky et al.’s study, where the phased and imputed genotypes in our study were directly obtained from [6]. Briefly, SHAPEIT2 [18] was used for phasing and IMPUTE2 [19] was used for imputation against the 1000 Genome reference panel containing 2, 504 individuals and ~82 million SNPs. Based on the phased and imputed SNPs, we extracted 6, 211, 048 of high confidence imputed SNPs (the ones with at least one heterozygote in the sample) in total.

#### Identification of imprinted genes and genes with dominance effect

We selected 22 known imprinted genes based on the list reported by Jadhav et al. [9]. These genes were selected because they had abundant expression in the 30 samples (S1 Table). For each potential imprinted gene, all SNPs in the gene coding region were selected as candidate *cis*-eQTLs. For each *cis*-eQTL and gene expression pair, the Stat-POE method was applied to detect candidate eQTLs with additive, dominance and POE effects. Four covariates were adjusted in the model including the total read counts per individual and the first three principal components computed from the matrix of normalized expression. In all of the hypothesis testing, Benjamini-Hochberg (BH) method was used for multiple comparison to adjust the p-values obtained from the LRT tests [20]. We tested the POE of the previously reported imprinted genes to evaluate the performance of our methods. For novel discovery of genetic effects of these imprinted genes, we tested the additive and dominance effects simultaneously.

## Acknowledgments

This work was supported by the internal support from University of South Carolina for Dr. Feifei Xiao. We are extremely grateful to Dr. Wei Sun and Dr. Zhabotynsky’s generous support who kindly provided the phased and imputed genotypes and pre-processed RNA-seq data.

## Supporting information

**S1 Table. List of imprinted genes identified in previous studies.** 22 genes were included to study the *cis*-eQTL and imprinting effects. These genes were included in Jadhav et al., 2018 which were compared to the identified imprinted genes reported as imprinted by the Genotype-Tissue Expression (GTEx) project (Baran et al. 2015). The genes were annotated by NCBI hg19 build. S1 File.

**S2 Table. A list of *cis*-eQTL (p-values < 0.05) for the 22 imprinting candidate genes.** ATL: alternative allele; a: additive effect; d: dominance effect; l: imprinting effect; expression: imprinting status. The p-values were adjusted by the BH multiple comparison method.

